# A subgroup of mitochondrial extracellular vesicles discovered in human melanoma tissues are detectable in patient blood

**DOI:** 10.1101/174193

**Authors:** Su Chul Jang, Rossella Crescitelli, Aleksander Cvjetkovic, Valerio Belgrano, Roger Olofsson Bagge, Johanna L. Höög, Karin Sundfeldt, Takahiro Ochiya, Raghu Kalluri, Jan Lötvall

## Abstract

Extracellular vesicles (EVs), including exosomes and microvesicles, are secreted from all cells, and convey messages between cells in health and disease. However, the diversity of EV subpopulations are only beginning to be explored. Since EVs have been implicated in tumor microenvironmental communication, we started to determine the diversity of EVs specifically in this tissue. To do this, we isolated EVs directly from patient melanoma metastatic tissues. Using EV membrane isolation and mass spectrometry analysis, we discovered enrichment of mitochondrial membrane proteins in the melanoma tissue-derived EVs, compared to non-melanoma-derived EVs. Specifically, EVs positive for a combination of the two mitochondrial inner membrane proteins MT-CO2 (mitochondrial genome) and COX6c (nuclear genome) were detected in the plasma of melanoma patients, and in ovarian and breast cancer patients. Furthermore, this subpopulation of EVs, contains active mitochondrial enzymes. Our findings show that tumor tissues are enriched in EVs with mitochondrial proteins and enzymatic activity, and these EVs can be detected in blood.

## Introduction

Extracellular vesicles (EVs) are nano-sized (50–1000 nm in diameter) vesicles with a lipid bi-layer membrane that play a significant role in mediating intercellular communication (1, 2). They can do this by either activating surface receptors of recipient cells or by transferring cargo proteins (3), nucleic acids (4), or lipids (5) to recipient cells. The interest in EVs has expanded exponentially over the last five to ten years due to their ability to mediate diverse biological effects, and shuttle molecules between cells. EVs can be found in all body fluids, including blood (6), urine (7), ejaculate (8), and breast milk (9), and they are considered to carry signatures of the cells that produce them. This means that EVs have promising potential as diagnostic markers in disease, especially for cancers. Examples of diagnostic EV markers that have been proposed in cancer to date include glypican-1 protein (10), EpCAM protein (11), KRAS-mutated DNA (12), oncogenic mRNA (13), and microRNAs (13). However, most EV-based biomarker candidates have initially been identified in cell culture-derived EVs and might not be valid markers for actual human disease. Here, we isolate and characterize EVs directly from human melanoma metastatic tissues, with the hypothesis that this is an environment enriched in tumor-associated EVs.

## Results

### Isolation and characterization of EVs directly from tumor tissues

First, the existence of EVs in the interstitial space of a melanoma metastatic tissue was confirmed by electron microscopy of tumor tissue pieces (Fig. 1*A*). Then, melanoma metastatic tissue-derived EVs were isolated using an ultracentrifugation-based protocol that is able to separate EV subpopulations; larger vesicles are isolated at lower speed (16,500 × *g*) and smaller vesicles at higher speed (118,000 × *g*). These EV subpopulations were further characterized by electron microscopy and RNA profile analysis, and showed typical morphology and RNA profiles of the EV subpopulations. Larger vesicles were 100–200 nm in diameter (Fig. 1*B*) and had an RNA profile comparable to microvesicles (MVs) (14) – a subpopulation of EVs that are considered to be produced by membrane budding – with the presence of 18S and 28S ribosomal RNA peaks and small RNAs (Fig. 1*D*). In contrast, smaller vesicles were 40–100 nm in diameter (Fig. 1*C*) and exhibited RNA profiles similar to exosomal RNA profiles (14), without prominent ribosomal RNA peaks (Fig. 1*E*). However, proteomics studies showed that majority of proteins are common in both larger and smaller vesicles, although some proteins are unique in each vesicle (15, 16). In addition, with current state of art isolation methods, each EV subpopulation are not possible to fully separate from each other. For these reasons, both larger and smaller vesicles were combined for further purification with an iodixanol density gradient, and subsequent proteomic experiments.

**Fig. 1.**
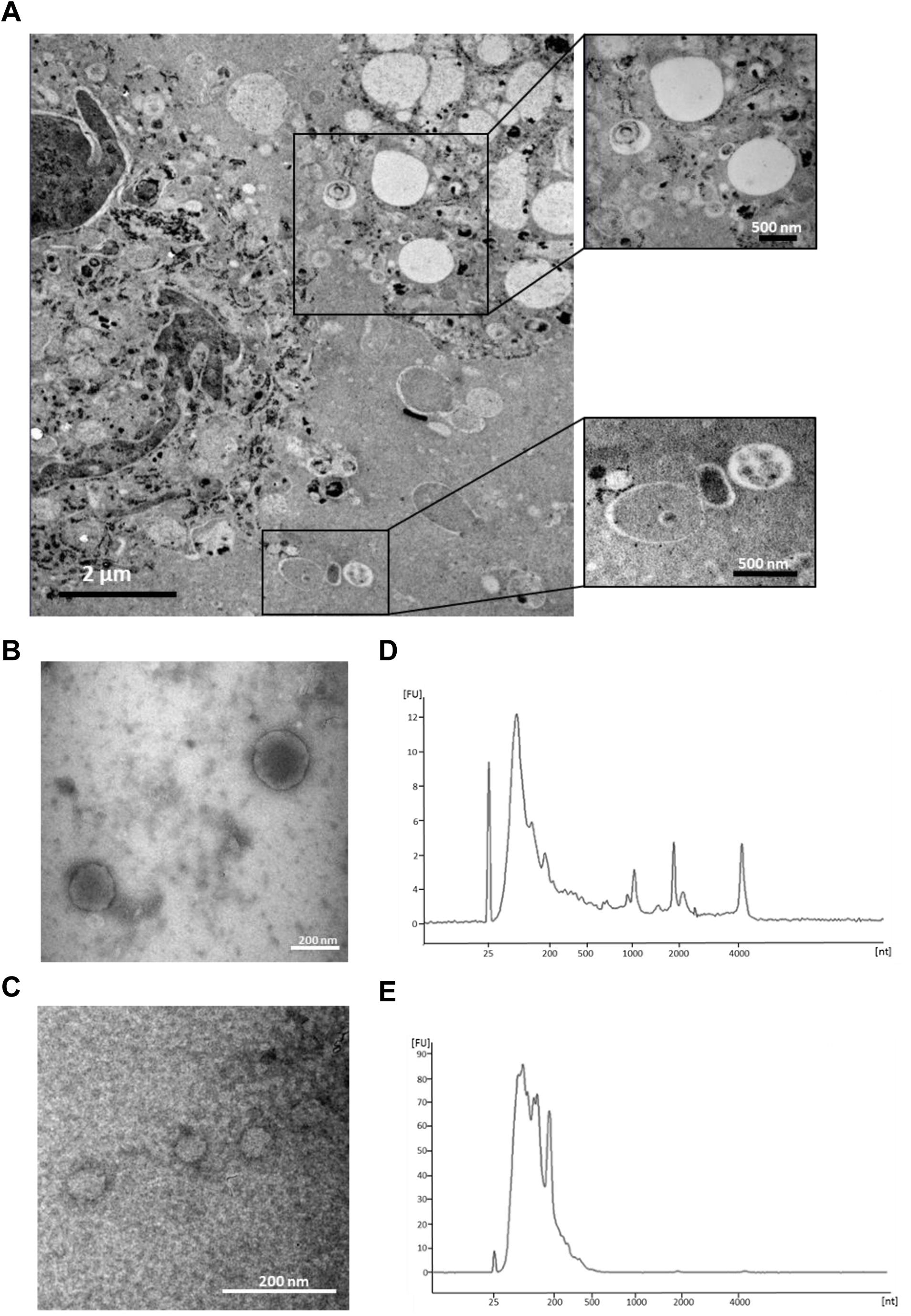
EVs in melanoma metastases tissue interstitial space were isolated and characterized. (*A*) Melanoma metastatic tissue consisted of cells containing melanin accumulations and EVs. Higher magnification pictures showed numerous different types of EVs in the tissue interstitial space. Electron microscope images of larger (*B*) and smaller (*D*) EVs. Larger EVs have a size range of 100–300 nm, and smaller EVs have a diameter of approximately 40–100 nm. RNA profiles of larger (*C*) and smaller (*E*) EVs. Larger EVs have prominent 18S and 28S ribosomal RNA peaks, whereas smaller EVs have small RNA with no or very small ribosomal RNA peaks.

### Identification of melanoma-enriched membrane proteins on EVs

To compare the surface protein expression, EVs that were isolated from five human melanoma metastatic tissues (referred to as MeT1 to MeT5), three cell lines (HEK293T, TF1, and HMC1), and one primary cell (mesenchymal stem cells, MSCs) were treated with high pH solution (200 mM sodium carbonate, pH 12) in order to open the membrane structure as described previously (17). The high pH-treated membranes were isolated with an iodixanol density gradient after a high salt (1 M KCl) treatment to remove proteins that are ionically bound to the membrane. After this treatment, removal of dark pigment from pigmented melanoma EVs was clearly observed (Fig. S1). The proteins in these membrane-enriched samples were analyzed by mass spectrometry. In total, 2592, 2862, 3461, 2239, 2750, 3577, 2096, 3521, and 1339 proteins were identified from MeT1-, MeT2-, MeT3-, MeT4-, MeT5-, HEK293T-, MSC-, TF1-, and HMC1-EVs, respectively. The identified proteins from different sources overlapped well with the EV proteome database EVpedia (18) (Fig. S2*A*). The overlapping percentages for MeT1-, MeT2-, MeT3-, MeT4-, MeT5-, HEK293T-, MSC-, TF1-, and HMC1-EVs were 74%, 79%, 77%, 84%, 76%, 79%, 94%, 83%, and 94%, respectively. These were all significantly higher than randomly selected proteins from the human proteome (38 ± 1.7%) (Fig.S2*B*). Similarly, 23.6% and 49.5% of proteins were uniquely found in EVs isolated from human and mice brain, respectively, compared with current EV proteome databases (19), suggesting that tissue-derived EVs were different from cell culture-derived EVs. In addition, classical EV marker proteins such as CD9, CD81, CD63, Syntenin-1, and Flotillins (20) were detected in all EVs with similar abundance, with few exceptions (Fig. S3). These results suggest that EVs from tumor tissues are indeed EVs rather than necrotic particles or artifacts of the isolation procedure.

Only membrane proteins that are annotated in the Uniprot database (21) were selected and compared to determine the common surface protein profiles of five different EV populations from melanoma metastatic tissues (Fig. 2*A*). Interestingly, membrane proteins from five MeT-EVs were distinct from those from non-melanoma-EVs (Fig. S4). In total, 1004 proteins that were identified in at least four MeT-EVs were selected for further analysis. In addition, 1422 membrane proteins that were identified in at least one of non-melanoma-derived EVs (HKE293T-, MSC-, TF1-, and HMC1-EVs) were selected and compared with membrane proteins from MeT-EVs to find unique membrane proteins on MeT-EVs. One hundred and thirty-six proteins were selected as melanoma-specific surface proteins, as they were not detected in the other EV membrane isolates (Fig. 2*B*). In addition, 582 proteins were selected among the 868 common proteins, because these proteins were 10-fold higher in abundance in the MeT-EVs compared to non-melanoma-derived EVs (Fig. 2*B*). To increase the confidence of the identification of melanoma specific molecules, the relative abundance of selected surface proteins was taken into account. Finally, 236 proteins that were highly abundant in all 5 MeT-EVs were selected as melanoma-specific surface molecules. (Dataset S1). Surprisingly, the percentage of endoplasmic reticulum and mitochondrial membrane proteins in candidates was higher than 25 percent (Fig. 2*C*). In general, the proportion of mitochondrial membrane proteins (6.9 ± 7.9%) were lower than plasma membranes (74.9 ± 13.2%) when the membrane proteins were analyzed from previous proteomic studies (43 datasets from EVpedia) (Fig. S5). Therefore, we focused on the mitochondrial membrane proteins for the further analysis. The relative abundance and the identification count on EVpedia of mitochondrial proteins were plotted (Fig. 2*D*). COX6c, SLC25A22, and MT-CO2 – all of which are mitochondrial inner membrane proteins – were selected for validation and tested for their presence in tumor tissue and cancer patient circulation, because they were relatively highly abundant and less commonly identified in other EV proteomic studies (Fig. 2*D*). These three proteins were highly expressed in melanoma metastatic tissue-derived EVs compared to non-melanoma-derived EVs (Fig. 2*E-G*). In addition to mitochondrial membrane proteins, HLA-DR (a plasma membrane protein) and Erlin2 (an endoplasmic reticulum membrane protein) were also highly expressed in melanoma metastatic tissue-derived EVs (Fig. S6).

**Fig. 2.**
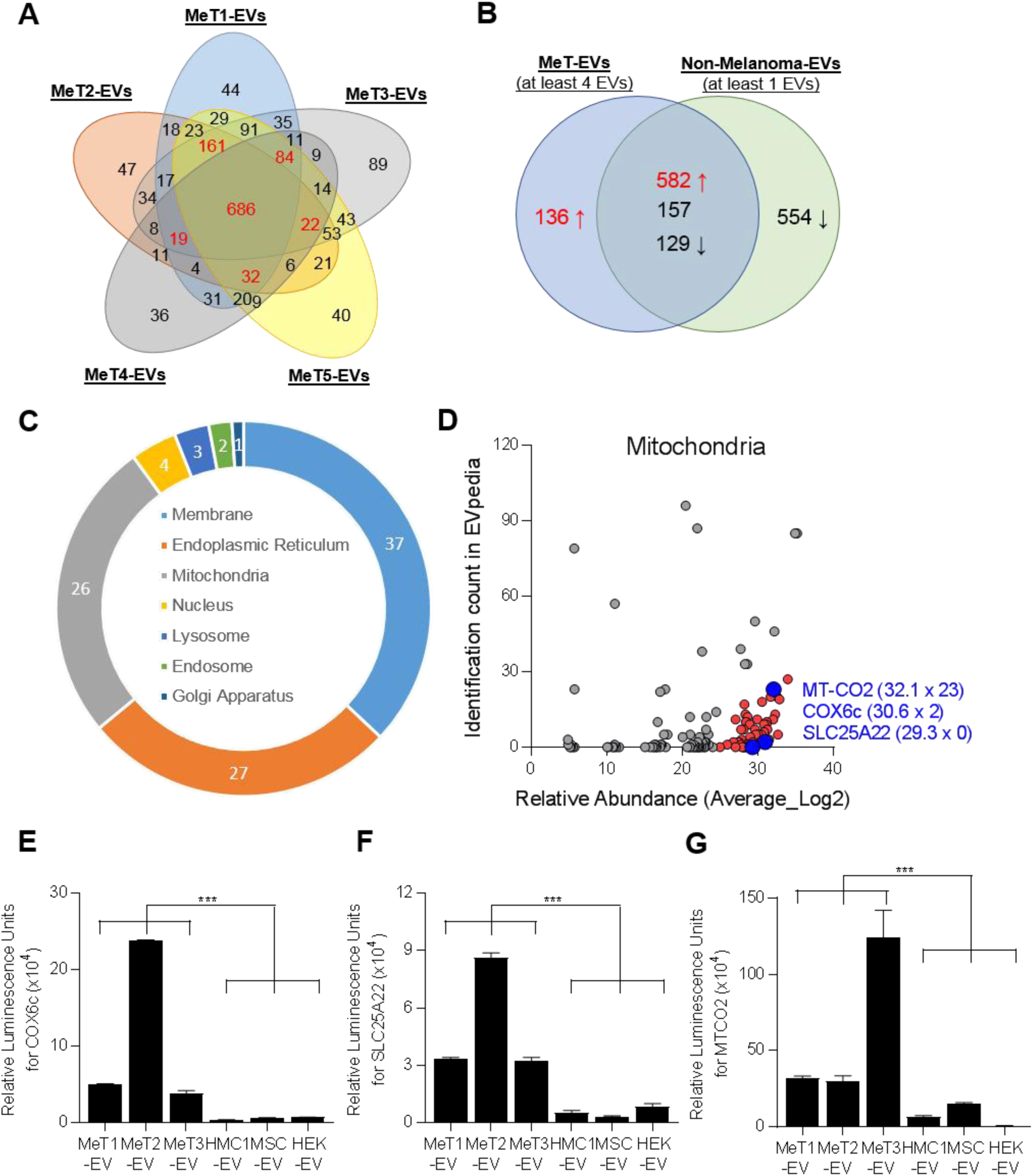
Proteomic analysis of EVs reveals the existence of mitochondrial membrane proteins. (*A*) Only membrane-localized proteins that were identified from five melanoma tissue-derived EVs (MeT1- to MeT5-EVs) were selected and compared. Numbers with a red color are proteins that were identified in at least 4 MeT-EVs. (*B*) Membrane proteins from MeT-EVs were compared with membrane proteins from non-melanoma-EVs. Common proteins were categorized by 10-fold difference of relative abundance. (*C*) The sub-cellular localization of 236 candidates was analyzed. (*D*) Mitochondrial membrane proteins were plotted with their relative abundance from a mass spectrometry analysis and their identification count from the EVpedia database. Blue color are the final three candidates for validation. IC; identification count. Three mitochondrial membrane proteins – COX6c (*E*), SLC25A22 (*F*), and MT-CO2 (*G*) – were experimentally validated with direct ELISA. Data are presented as the mean ± SD. \*\*\**p* < 0.001.

### Existence of mitochondrial proteins on exosomes

It is known that cancer cells are metabolically active and have enhanced mitochondrial biogenesis (22). However, mitochondrial genomes are reduced in many cancers, which is considered to support tumor progression (23, 24). Furthermore, whole mitochondria can be transferred between cells in disease conditions such as cancer (25), stroke (26), and lung injury (27), but the details on the mechanism of transfer remains elusive. Recently, MVs from MSCs, a subpopulation of EVs secreted by membrane budding, were reported to contain some mitochondrial components, including proteins and mtDNA (28). These findings suggest that mitochondrial components are secreted from the cells in the form of EVs. However, it is still unknown whether mitochondrial proteins are secreted as exosomes (EXOs), a subpopulation of EVs secreted by an endocytic pathway. To test this, EXOs were isolated from the MML1 and HMC1 cell lines by differential centrifugation coupled with an iodixanol density gradient. The mitochondrial protein MT-CO2, which is translated from the mitochondrial genome (29), was detected in the same density gradient fractions where the exosomal marker CD81 (Fig. S7 *A* and *B*) and most of the EVs are present (Fig. S7 *C* and *D*). Surface expression of MT-CO2 as well as the exosomal markers CD9 and CD81 was detected in EXOs isolated from MML1 cells (MML1-EXOs) (Fig. S7*E*) and HMC1 cells (HMC1-EXOs) (Fig. S7*F*). However, its expression was higher in MML1-EXOs compared with HMC1-EXOs (Fig. S7*G*), which was consistent with our initial result (Fig. 2*G*).

EXOs that contain MT-CO2 (MTCO2-EXOs) were further isolated from EV isolates, and their protein profile was determined by mass spectrometry. EXOs that contain FACL4 (FACL4-EXOs), another mitochondrial membrane protein but translated from the nuclear genome, were used as an additional control. In total, 449, 646, and 839 proteins were identified from EXOs, FACL4-EXOs, and MTCO2-EXOs, respectively (Dataset S2). Venn diagram analysis showed that 410 proteins were common for all three groups, and 179 proteins were uniquely identified in MTCO2-EXOs (Fig. 3*A*). Heatmap analysis based on the relative abundance revealed that each sample has a distinct proteome profile (Fig. 3*B*), and these proteins were categorized into the following five clusters: ‘Common’, ‘Mito-EXOs excluded’, ‘FACL4-EXOs enriched’, ‘FACL4/MTCO2-EXOs enriched’, and ‘MTCO2-EXOs enriched’. Importantly, most of the classical marker proteins such as tetraspannins, TSG101, Alix, Annexins, and Flotillins were present in all three EXO populations (Dataset S3), implying that there may be a crosstalk between mitochondria and the exosome biogenesis pathways. Further, gene ontology analysis showed that FACL4-EXOs and MTCO2-EXOs clusters were enriched with proteins involved in metabolic processes (Fig. 3*C*). In addition, protein-protein interaction networks showed that mitochondrial proteins, including ATP synthase subunits in the EVs were interrelated to each other (Fig. S8), and the activity of ATP synthase was higher in MTCO2-EXOs compared to EXOs (Fig. 3*D*). These results suggest that subpopulations of EVs, including both MVs and EXOs, contain active forms of mitochondrial proteins.

**Fig. 3.**
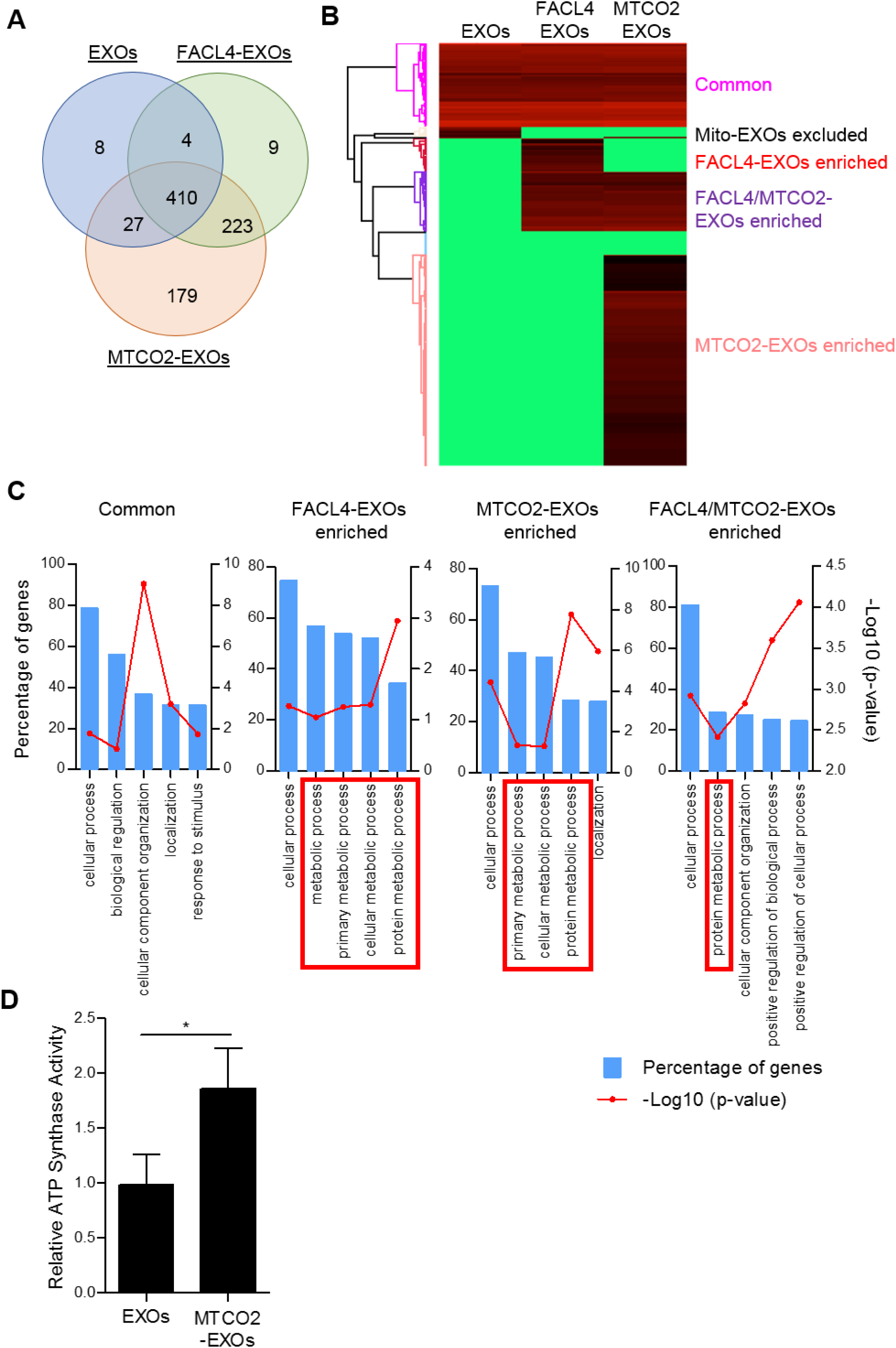
Subpopulations of EXOs harbor mitochondrial proteins. Subpopulations of EXOs were isolated with antibodies against FACL4 or MT-CO2, and their proteomes were analyzed. The identified proteins were compared by Venn diagram (*A*) and heatmap analysis (*B*). Based on relative abundance of proteins, five different clusters were identified (*B*), and gene ontology analysis was performed for 4 of the 5 clusters (*C*). FACL4-EXOs and/or MTCO2-EXOs clusters enriched with metabolic process-related proteins are shown in red boxes. (*D*) ATP synthase activity was measured for EXOs and MTCO2-EXOs. Data are presented as the mean ± SD. \**p* < 0.05.

### Detection of mitochondrial extracellular vesicles in patient plasma

We have shown that mitochondrial proteins are present on the surface of the subpopulations of EVs and are more abundant in melanoma-derived EVs. Finally, to test whether EVs with mitochondrial membrane proteins leak out to the circulation of cancer patients, we developed a sandwich ELISA detection system with MT-CO2 and COX6c antibodies (Fig. 4*A*). This sandwich ELISA was first tested with purified EVs from melanoma tissues and cell lines. High luminescence signals were observed only in melanoma tissue-derived EVs (Fig. 4*B*), which is in line with our proteomics and direct ELISA results (Fig. 2*E-G*). Because both MT-CO2 and COX6c are membrane proteins, we hypothesized that the luminescent signal seen in our sandwich ELISA system is indeed associated with membrane structures such as EVs. To further test this hypothesis, we measured the presence of these markers directly in plasma from melanoma patients without any preceding EV isolation. We detected significantly higher levels of combined MT-CO2 and COX6c in plasma of melanoma patients compared with healthy controls (*p* = 0.0038) (Fig. 4*C*). To test whether this increased concentration of MT-CO2/COX6c positive EVs is specific for melanoma, we applied the same assay to plasma from ovarian and breast cancer patients. To our surprise, the concentration of MT-CO2/COX6c was increased in the plasma of both patients with ovarian (*p* = 0.0220) (Fig. 4*D*) and breast cancer (*p* = 0.0408) (Fig. 4*E*), with significant differences *vs* healthy controls observed. However, there were no significant differences in the concentration of these EVs between patients with malignant or benign ovarian cysts, because those with the benign disease also had significant increase of MT-CO2/COX6c EVs (data not shown). Overall, these findings suggest that mitochondrial proteins on EVs are released in several malignant diseases of different cellular origin, including melanoma, ovarian cancer, and breast cancer. Further exploration of the presence of mitochondria-derived EVs in other types of tumors, both malignant and benign, is warranted.

**Fig. 4.**
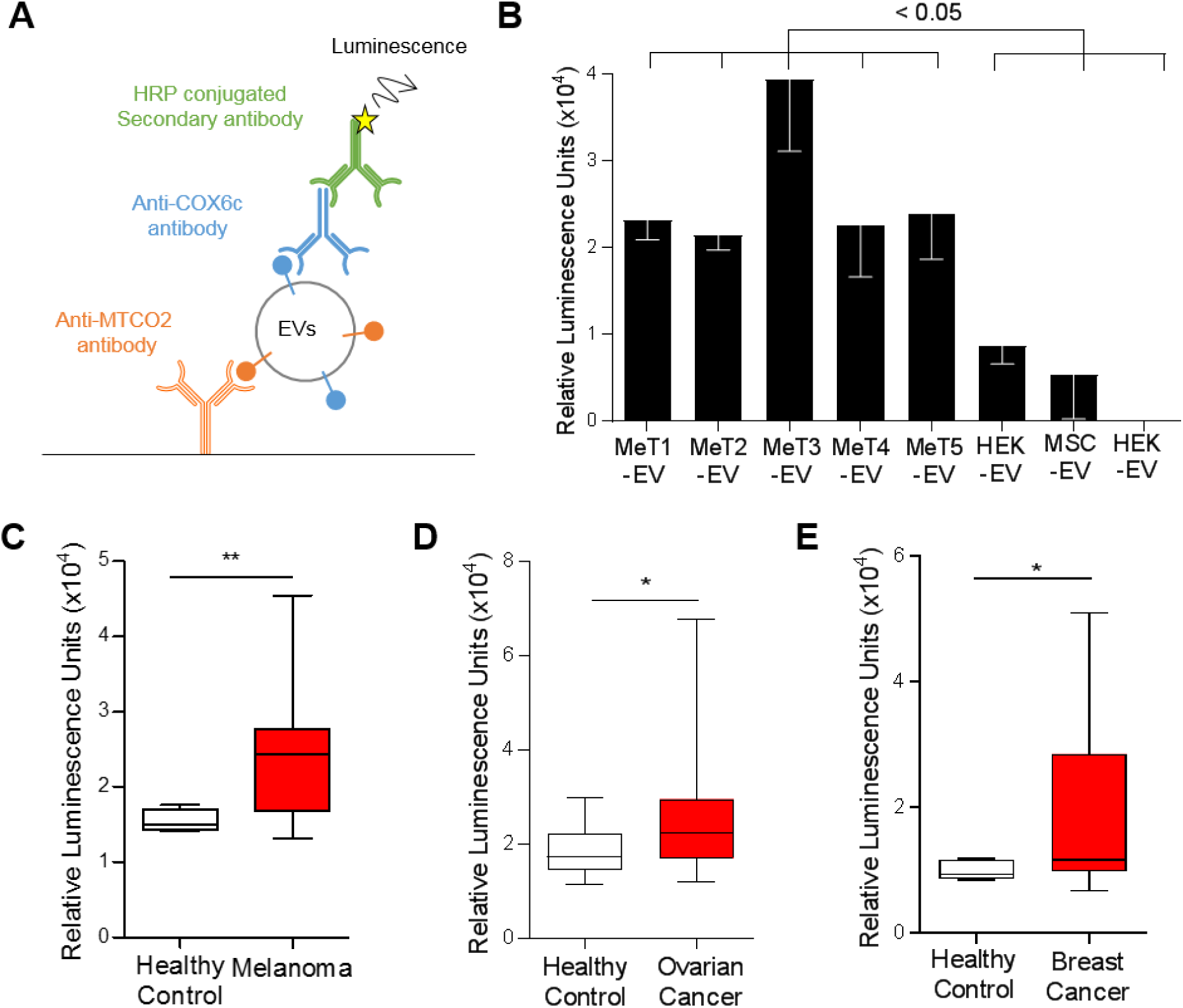
Mitochondrial membrane proteins on the surface of EVs are unique biomarkers for cancer. (*A*) Schematic illustration of the sandwich ELISA. MT-CO2 antibody was coated on the plate to capture EVs, and COX6c antibody was used to detect the EVs. (*B*) The sandwich ELISA system was validated with melanoma tissue-derived EVs and non-melanoma-derived EVs. Data are presented as the mean ± SD. The levels of mitochondrial proteins (MT-CO2 and COX6c) were examined in blood plasma from melanoma patients (*n* = 26) (*C*), ovarian cancer patients (*n* = 62) (*D*), and breast cancer patients (*n* = 13) (*E*). Healthy control was six. Whiskers show the minimum-maximum and lines inside box represent the median. \**p* < 0.05, \*\**p* < 0.001.

## Discussion

Liquid biopsy is a powerful method to detect or monitor disease states, especially for cancer, from body fluids such as blood or urine, as it is giving noninvasiveness and better snapshot of heterogeneity of tumor tissues (30). Recently, EVs have been focused for advantageous materials over the current circulating cells and DNA. EVs modulate the tumor progression (31), metastasis (32), and angiogenesis (5) by shuttling their cargo that reflect their parental cell status. Those EVs are secreted into the interstitial space of tissues, and sometimes can leak and accumulate in the circulation, which makes them potential circulating biomarkers. Numerous studies have been attempted to find EV-based protein biomarkers from body fluids of patients including blood or urine. However, there are many of non-cancer-derived EVs in circulation and they may have hindered successful identification of markers. Here, we hypothesized that tumor interstitial space is full of tumor-derived EVs. Isolating EVs from tissues has been reported from mouse brain (33), but our study is the first study isolating EVs from human patient tumor tissues. In addition, we have applied the membrane proteomics to identify the surface markers. Membrane proteins have been focused intensively since they are entry or exit of cellular signals, which making them important druggable targets as well as surface markers (34). Although EVs are known to be enriched with membrane proteins, more than 60% of identified proteins are non-membrane proteins according to EVpedia (18). By applying membrane proteomics, many of membrane proteins were newly identified from our study.

Interestingly, our study describes that melanoma tissue-derived EVs were enriched with mitochondrial membrane proteins, compared with non-melanoma-derived EVs. Furthermore, our study shows that subpopulation of EXOs contain mitochondrial components and have a distinct protein profile. The release of mitochondrial components to the extracellular space could be related with enhanced mitochondrial function in cancer, since mitochondria is closely connected with cancer in terms of progression, signaling, and survival (23, 24). It has been discovered that mitochondria secret the vesicles inside of cells. The fate of these vesicles is either fusing with lysosome to undergone a degradation pathway (35) or fusing with ER-derived pre-peroxisomes to generate the peroxisomes (36). However, secretion to extracellular space or fusion with exosomal pathway of those mitochondrial vesicles has not been described. This is interesting because current state of art of EV biogenesis does not elucidate the connection with mitochondria. Elucidating the in-depth mechanism how mitochondria and EV biogenesis pathway are interconnected is required in future to fully understand biogenesis of EVs.

We have found not only mitochondrial membrane proteins, but also many of other membrane proteins, from different cellular compartments including plasma membrane and endoplasmic reticulum membrane, that could be another good biomarker candidates. For example, HLA-DR, which is a plasma membrane protein and was in our biomarker candidates, was used to predict response to anti-PD1/PD-L1 therapy by detecting HLA-DR on melanoma tissue (37). In addition, an endoplasmic reticulum membrane protein, Erlin-2, is related with survival of breast cancer by modulating endoplasmic reticulum stress pathways (38). The diagnostic potential of these proteins is needed to verify in future.

In summary, we have isolated EVs from melanoma metastatic tissues from human patients and analyzed their surface proteomes, detecting important differences between tissue-derived EVs and classical cell line-derived EVs. Most importantly, mitochondrial membrane proteins were present at high levels in melanoma tissue-derived EVs, and can be detected in increased concentrations in the plasma of melanoma patients, but also in patients with ovarian- or breast cancer. In addition, we found that certain subpopulations of EXOs harbor both active mitochondrial proteins and classical EV markers, which is a new paradigm in EV biology.

## Materials and Methods

### Human samples

Tissues from melanoma lymph node or skin metastases were obtained from patients during surgery and were preserved in complete cell media (without fetal bovine serum, FBS) at 4°C and were used to isolate EVs. A total of 20 ml of peripheral blood was collected from melanoma patients, breast cancer patients, and healthy controls in EDTA tubes. Plasma was obtained by centrifugation at 1880□ × □*g* for 10 min, followed by a second centrifugation at 2500□ × □*g* for 10 min. All centrifugations were performed at 4°C. The study was approved by the Regional Ethical Review Board at the University of Gothenburg (096-12), and all participants provided a written informed consent. Ovarian plasma was chosen from a previously described cohort collected between 2001 and 2010 (39). Blood samples were collected after anesthesia bur before surgery. Six ml of blood were collected in EDTA vacutainers using standardized procedures, centrifuged and directly aliquoted into Eppendorf tubes, frozen and stored at -80°C within 30-60 minutes after withdrawal. The selected samples had seen one freeze thaw cycle only. Ethical board approval number are 348-02 and 201-15.

### Cell culture

The human mast cell line HMC1 and TF1 cells were cultured in IMDM (HyClone, Logan, UT). HEK293T and MML1 cells were cultured in RPMI1640 (Sigma Aldrich, St Louis, MO) media. Media was supplemented with 10% EV-depleted FBS (Sigma Aldrich), 2 mM L-glutamine (HyClone), and 1.2 U/ml 1-thioglycerol (Sigma Aldrich). The human MSCs from bone marrow were obtained as passage 1 from the MSC distribution of the Institute of Regenerative Medicine at Scott and White, USA, and cultured in alpha minimum essential medium (GIBCO^®^ GlutaMAX^™^, Invitrogen, Carlsbad, CA) supplemented with 15% EV-depleted FBS (Sigma Aldrich). Three to four passages of MSCs were used for EV isolation. All media contained 100 U/ml penicillin and 100 μg/ml streptomycin (HyClone). For the EV depletion, FBS was ultracentrifuged at 118,000 × *g*_avg_ (Type 45 Ti rotor, Beckman Coulter, Miami, FL) for 18 h and filtered through a 0.22 μm filter as previously described (40).

### Isolation of EVs from melanoma metastases tissue

Subpopulations of EVs were isolated from melanoma metastases using a centrifugation-based protocol. Tumor pieces were gently sliced into small fragments (1–2 mm) and incubated with collagenase D (Roche, Basel, Switzerland) (2 mg/ml) and DNase I (Roche) (40 U/ml) dissolved in RPMI plain medium (Sigma Aldrich) for 30 min at 37°C. After a filtration step (70 μm pore size), cells and tissue debris were eliminated by centrifugation at 300 × *g* for 10 min and 2000 × *g* for 20 min. Supernatants were centrifuged at 16,500 × *g*_avg_ (Type 45 Ti) for 20 min and 110,000 × *g*_avg_ (Type 45 Ti) for 2.5 h to collect larger vesicles and smaller vesicles, respectively. All centrifugations were performed at 4°C. Pellets were resuspended in PBS. Larger and smaller vesicles were combined and further purified by an isopycnic centrifugation using an iodixanol gradient (OptiPrep^TM^, Sigma-Aldrich).

### Isolation of EVs from cell lines

Conditioned media from cell cultures was harvested and centrifuged at 300 × *g* for 10 min to remove cells. The supernatant was then centrifuged at 2,000 × *g*_avg_ for 20 min to remove apoptotic bodies and cell debris. MVs and EXOs were pelleted at 16,500 × *g*_avg_ (Type 45 Ti) for 20 min and at 118,000 × *g*_avg_ (Type 45 Ti) for 3.5 h. Pellets were resuspended in PBS. MVs and EXOs pellets were combined and further purified by an isopycnic centrifugation using an iodixanol gradient.

### Iodixanol density gradient

EVs (MVs and EXOs) in PBS (1 ml) were mixed with 60% iodixanol (3 ml) and laid on the bottom of an ultracentrifuge tube followed by addition of 30% iodixanol (4 ml) and then 10% iodixanol (3 ml). Samples were ultracentrifuged at 178,000 × *g*_avg_ (SW 41 Ti, Beckman Coulter) for 2 h. EVs were collected from the interface between the 30% and 10% iodixanol layers. For the EXOs purification, pelleted EXOs (1 ml) were mixed with 60% iodixanol (3 ml) and laid on the bottom of an ultracentrifuge tube. A discontinuous iodixanol gradient (35%, 30%, 28%, 26%, 24%, 22%, and 20%; 1 ml each, but 2 ml for 22%) was overlaid. Samples were ultracentrifuged at 178,000 × *g*_avg_ (SW 41 Ti) for 16 h. EXOs were collected from the interface of the 20% and 22% iodixanol layers. Collected EVs or EXOs were diluted with PBS (up to 94 ml) and ultrancetrifuged at 118,000 × *g*_avg_ (Type 45 Ti) for 3.5 h. The pelleted EVs and EXOs were resuspended in PBS.

### Membrane isolation

Isolated EVs were incubated with 100 mM sodium carbonate solution (pH 12) for 1 h at room temperature with rotation. Potassium chloride solution (1 M) was added and further incubated for 1 h. Samples were subjected to iodixanol density gradient purification as described above for EVs, and membranes were collected from the interface between the 30% and 10% iodixanol layers.

### Transmission electron microscopy

One melanoma metastatic tissue from lymph node was acquired during surgery, and the sample was dissected into small pieces. Samples were placed in 150 μm deep membrane carriers (Leica Microsystems, Bensheim, Germany) that was filled with 20% BSA in PBS and high pressure frozen using an EMPactI (Leica Microsystems). A rapid freeze substitution protocol using 2% uranyl acetate (from a 20% methanolic stock solution) in dehydrated acetone for 1 h was then applied (41, 42). The temperature was increased by 3°C/hour to -50°C where samples stayed for the remainder of the protocol, including polymerization. Samples were washed two times with dehydrated acetone before starting infiltration with increasing concentrations of HM20 (3:1, 2:1, 1:1, 1:2, 1:3 acetone:HM20 for 2-3 hours each), followed with 3 changes with HM20 (2 h each and once overnight). Samples were polymerized in UV light for 48 h. Thin sections (70 nm) were cut and contrasted with 2% uranyl acetate in 25% ethanol (4 min) and Reynold’s lead citrate (2 min). For analysis of EVs, a drop (10 μl) of isolated EVs was placed on 200-mesh formvar/carbon copper grids (Ted Pella, Redding, CA) for 5 min and stained with UranyLess (EMS, Fort Washington, PA) for 1 min. Images were obtained using a LEO 912AB Omega 120 kV electron microscope (Carl Zeiss SMT, Mainz, Germany). Digital image files were acquired with a Veleta CCD camera (Olympus-SiS, Münster, Germany).

### RNA isolation and detection

RNA was isolated from melanoma metastases-derived EVs (larger and smaller vesicles) using the miRCURY^TM^ RNA Isolation Kit (Exiqon, Vedbaek, Denmark) according to the manufacturer’s protocol. EV RNA profiles were analyzed using capillary electrophoresis (Agilent RNA 6000 Nano Kit on an Agilent 2100 Bioanalyzer, Agilent Technologies, Palo Alto, CA). A total of 1 μl of RNA was analyzed according to the manufacturer’s protocol as previously described (43).

### Western blot analysis

Proteins from the iodixanol density gradients were separated by SDS-PAGE and transferred to a polyvinylidene fluoride membrane. The membrane was blocked with 5% non-fat dry milk for 2 h and then incubated with anti-CD81 (Santa Cruz Biotechnology, Santa Cruz, CA, sc9158) or anti-MT-CO2 (Abcam, Cambridge, UK, ab91317) at 4°C overnight. After secondary antibody incubation, the immune-reactive signals were visualized using SuperSignal™ West Femto Maximum Sensitivity Substrate (Thermo Fisher Scientific, San Jose, CA) with a VersaDoc 4000 MP (Bio-Rad Laboratories, Richmond, CA).

### Particle measurement

The numbers of particles from the iodixanol density gradients were measured using ZetaView^®^ PMX110 (Particle Metrix, Diessen, Germany). The chamber temperature was automatically measured and integrated into the calculation, and the sensitivity of the camera was set to 80. Data were analyzed using the ZetaView^®^ analysis software version 8.2.30.1 with a minimum size of 5, a maximum size of 5000, and a minimum brightness of 20. Samples were measured in triplicate.

### ELISA

For the direct ELISA, EVs were coated on 96-well plates overnight at 4°C. Plates were blocked with 1% BSA in PBS for 1 h and incubated with anti-CD9 (BD Biosciences, San Jose, CA, 555370), anti-CD81, anti-COX6c (Santa Cruz Biotechnology, sc390414), anti-SLC25A22 (Thermo Fisher Scientific, PA5-29181), or anti-MT-CO2 antibodies for 2 h. After washing, the appropriate secondary antibodies with HRP were added. The reaction was initiated by adding TMB substrate solution, terminated by 2M H_2_SO_4_, and the optical density was measured at a wavelength of 450 nm. For the sandwich ELISA, MT-CO2 antibody was coated on black 96-well plates overnight at 4°C. The MT-CO2 antibody was purified on a protein G column prior to use to remove the carrier proteins. Plates were blocked with 1% BSA in PBS for 1 h. EVs or body fluids were added to the wells and incubated for 2 h at room temperature. A total of 50 μl of blood plasma from patients was used. After washing, COX6c antibody was incubated for 1 h and then HRP-conjugated anti-mouse antibody was incubated for 1 h. Luminescent signal was obtained with the BM Chemiluminescence ELISA Substrate (BD Biosciences).

### Isolation of EXOs containing MT-CO2 or FACL4

Antibodies against MT-CO2 or FACL4 (Abcam, ab155282) were conjugated to magnetic beads with the Dynabeads^®^ Antibody Coupling Kit (Thermo Fisher Scientific) following the manufacturer’s instructions. Iodxianol-purified EXOs were incubated with anti-MT-CO2 or anti-FACL4 antibody-conjugated magnetic beads for 2 h at room temperature with rotation. Unbound EXOs were removed and washed with PBS. Bound EXOs were eluted with acidic washing buffer (10 mM HEPES, 10 mM 2-(N-morpholino) ethanesulfonic acid, 120 mM NaCl, 0.5 mM MgCl_2_, 0.9 mM CaCl_2_, pH 5) for 10 min.

### LC-MS/MS and protein search

The protein samples were digested with trypsin using the filter-aided sample preparation (FASP) method. Briefly, protein samples were reduced with 100 mM dithiothreitol at 60°C for 30 min, transferred onto 30 kDa MWCO Nanosep centrifugal filters (Pall Life Sciences, Ann Arbor, MI), washed with 8M urea solution and alkylated with 10 mM methyl methanethiosulfonate in 50 mM TEAB and 1% sodium deoxycholate. Digestion was performed in 50 mM TEAB, 1% sodium deoxycholate at 37°C in two stages: the samples were incubated with 300 ng of Pierce MS-grade trypsin (Thermo Scientific) for 3h, and then with 300 ng additional trypsin overnight. The digested peptides were desalted using Pierce C-18 spin columns (Thermo Scientific), the solvent was evaporated, and the peptide samples were resolved in 3% acetonitrile, 0.1% formic acid solution for LC-MS/MS analysis. Each sample was analyzed on a Q Exactive mass spectrometer (Thermo Fisher Scientific) interfaced with Easy-nLC 1200 nanoflow liquid chromatography system. Peptides were trapped on the C18 trap column (200 μm X 3 cm, particle size 3 μm), and separated on the home-packed C18 analytical column (75 μm X 30 cm, particle size 3 μm). A gradient from 8% to 24% B over 75 min, from 24% to 80% B over 5 min, and 80% for 10 min (solvent A: 0.2% formic acid, solvent B: 98% acetonitrile, 0.2% formic acid) was used at a flow rate of 200 nl/min. Precursor ion mass spectra were recorded in positive ion mode at a resolution of 70 000 and a mass range of 400 to 1600 m/z. The 10 most intense precursor ions were fragmented using HCD at a collision energy of 30, and MS/MS spectra were recorded in a scan range of 200 to 2000 m/z and a resolution of 35 000. Charge states 2 to 6 were selected for fragmentation, and dynamic exclusion was set to 30 s. Exclusion lists of m/z values of the identified peptides at 1% FDR with a 10 min retention time window were generated from each results file. Subsequently, a second LC-MS/MS analysis was performed using the same settings as before, besides the exclusion of m/z values present in the first LC-MS/MS analysis. The MaxQuant quantification tool with the Andromeda search engine (version 1.5.2.8) was used for the identification and quantification of proteins (44). Proteins were searched with the following parameters: enzyme specificity, trypsin; variable modification, oxidation of methionine (15.995 Da); fixed modification, carbamidomethylation of cysteine (57.021 Da); two missed cleavages; 20 ppm for precursor ions tolerance and 4.5 ppm for fragment ions tolerance; *Homo sapiens* reference proteome data from Swiss-Prot (20,196 entries); 1% false discovery rate; and a minimum peptide length of seven amino acids. The first major protein identified was chosen as the representative protein of each protein group and was used for further analysis. Normalized label-free quantification intensity of proteins was obtained by a label-free quantification tool, which is implemented in the MaxQuant software, with a minimum of two ratio counts.

### Systematic analysis

Protein localization was obtained from the Uniprot database and only primary localization of proteins was used for the analysis. Biological process terms from Gene Ontology were analyzed using DAVID (https://david.ncifcrf.gov/). The protein-protein interaction network and identification count were obtained from the STRING database (http://string-db.org/) and the EVpedia database (https://evpedia.info), respectively. Heat map was analyzed with Perseus software (http://www.coxdocs.org/doku.php?id=perseus:start).

### ATP synthase activity measurement

The activity of ATP synthase was measured using 10 μg of EXOs and MTCO2-EXOs with the ATP Synthase Enzyme Activity Microplate Assay Kit (Abcam) following the manufacturer’s instructions.

### Statistical analysis

GraphPad Prism software version 5 (GraphPad Software) was used for *p*-value calculation. The unpaired two-tailed Student’s *t*-test and one-way ANOVA with Turkey’s multiple comparison test were conducted for comparison between two samples and multiple samples, respectively. The unpaired two-tailed Student’s *t*-test with Welch’s correction was conducted for plasma samples. A *p*-value < 0.05 was considered to be significant.

## Acknowledgements

We thank Evelin Berger, Annika Thorsell, and Carina Sihlbom at the Proteomics Core Facility, University of Gothenburg, Sweden, for mass spectrometry analysis. The Proteomics Core Facility at Sahlgrenska Academy is grateful to the Inga-Britt and Arne Lundbergs Forskningsstiftlese for the donation of the Orbitrap Fusion Tribrid MS instrument. We thank Bengt R. Johansson for helpful discussions. We also thank Electron Microscopy Unit (EMU), and later, Centre for Cellular Imaging at the University of Gothenburg and the National Microscopy Infrastructure, NMI (VR-RFI 2016-00968) for providing assistance in microscopy. We thank Gunnar Nilsson at the Karolinska Institute, Stockholm, Sweden, for the kind gift of the human mast cell line HMC-1. This work was funded by the Swedish Research Council (K2014-85X-22504-01-3), VBG Group Herman Krefting Foundation for Asthma and Allergy Research, the Swedish Heart and Lung Foundation (20120528), the Swedish Cancer Foundation (CAN2014/844), and Basic Science Research Program through the National Research Foundation of Korea (NRF) funded by the Ministry of Education (2016R1A6A3A03007377).

## Competing interests

J.L. and S.C.J. are currently chief scientist and scientist, respectively, at Codiak Biosciences in Cambridge, MA, developing exosomes as therapeutics. S.C.J., R.C. and J.L. are developing a patent exploring EV biomarkers, and this patent is not related to Codiak Biosciences.

## Author Contributions

S.C.J. and J.L. designed the study. S.C.J. performed most of experiments and analyzed the data. C.R. and A.C. isolated EVs from melanoma tissues. C.R. and J.H. performed electron microscopy. R.O.B. and V.B. provided blood of melanoma patients and metastatic melanoma tissues. K.S. provided cystic fluids and blood from ovarian patients. The manuscript was written by S.C.J., C.R., and J.L. with the help of J.H., V.B., R.O.B., K.S, O.T., and R.K.

---

**Fig. S1.**
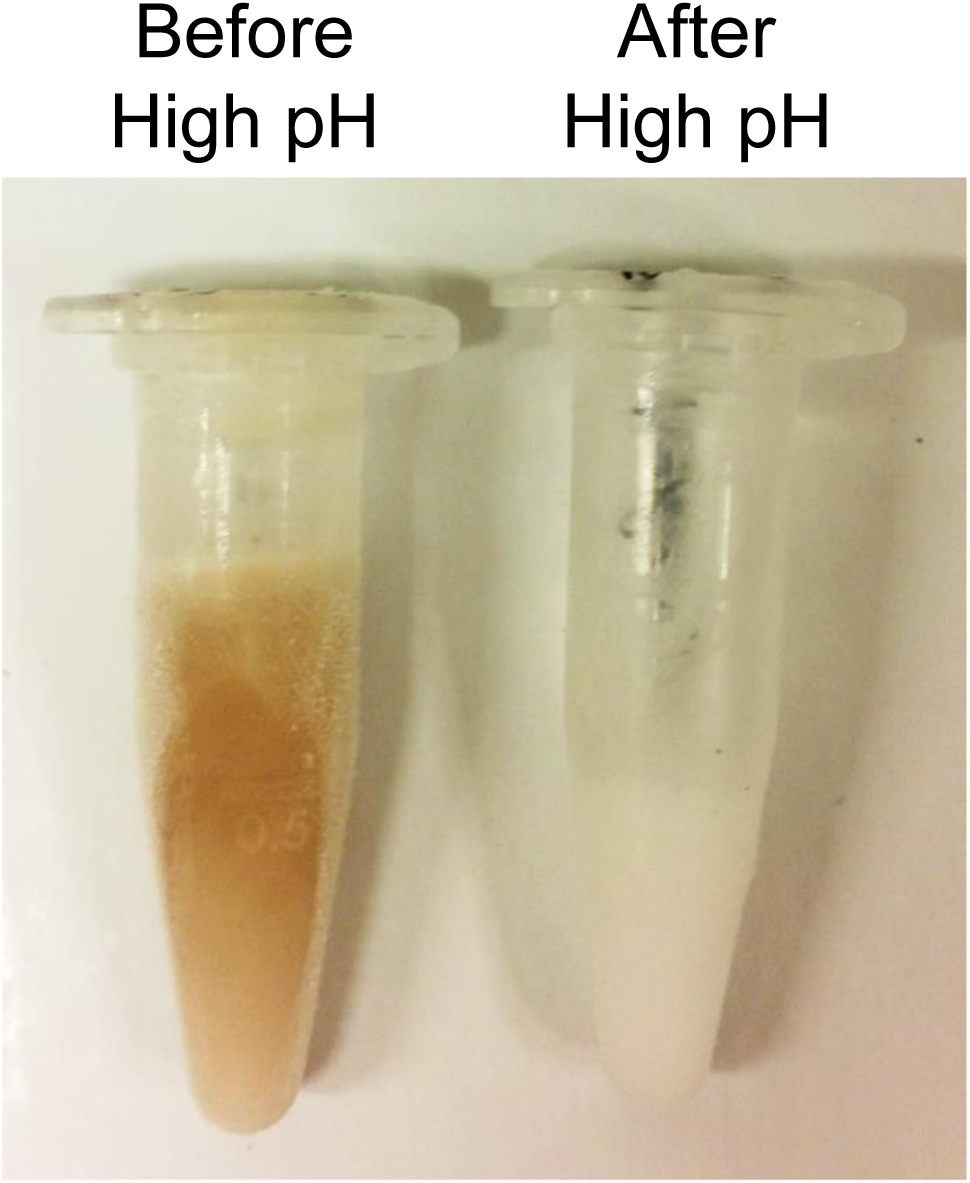
Image of EVs before and after high pH treatment. Brown color of melanin on EVs were clearly disappeared after high pH treatment.

**Fig. S2.**
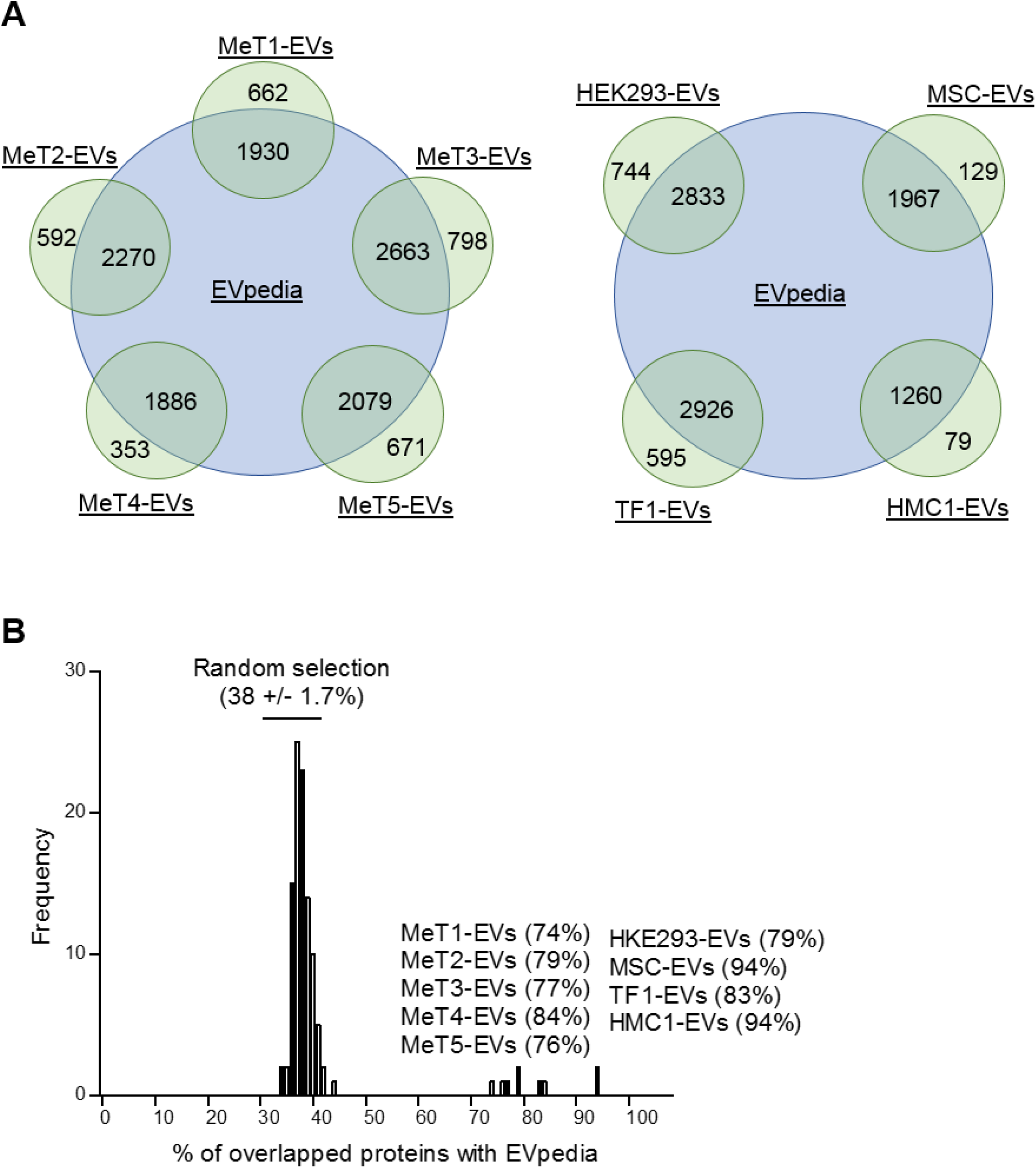
Comparison with an EV database. (*A*) Identified EV proteins were well overlapped with the EVpedia. (*B*) The overlapping percentage with EVpedia was higher in EVs compared to randomly selected proteins.

**Fig. S3.**
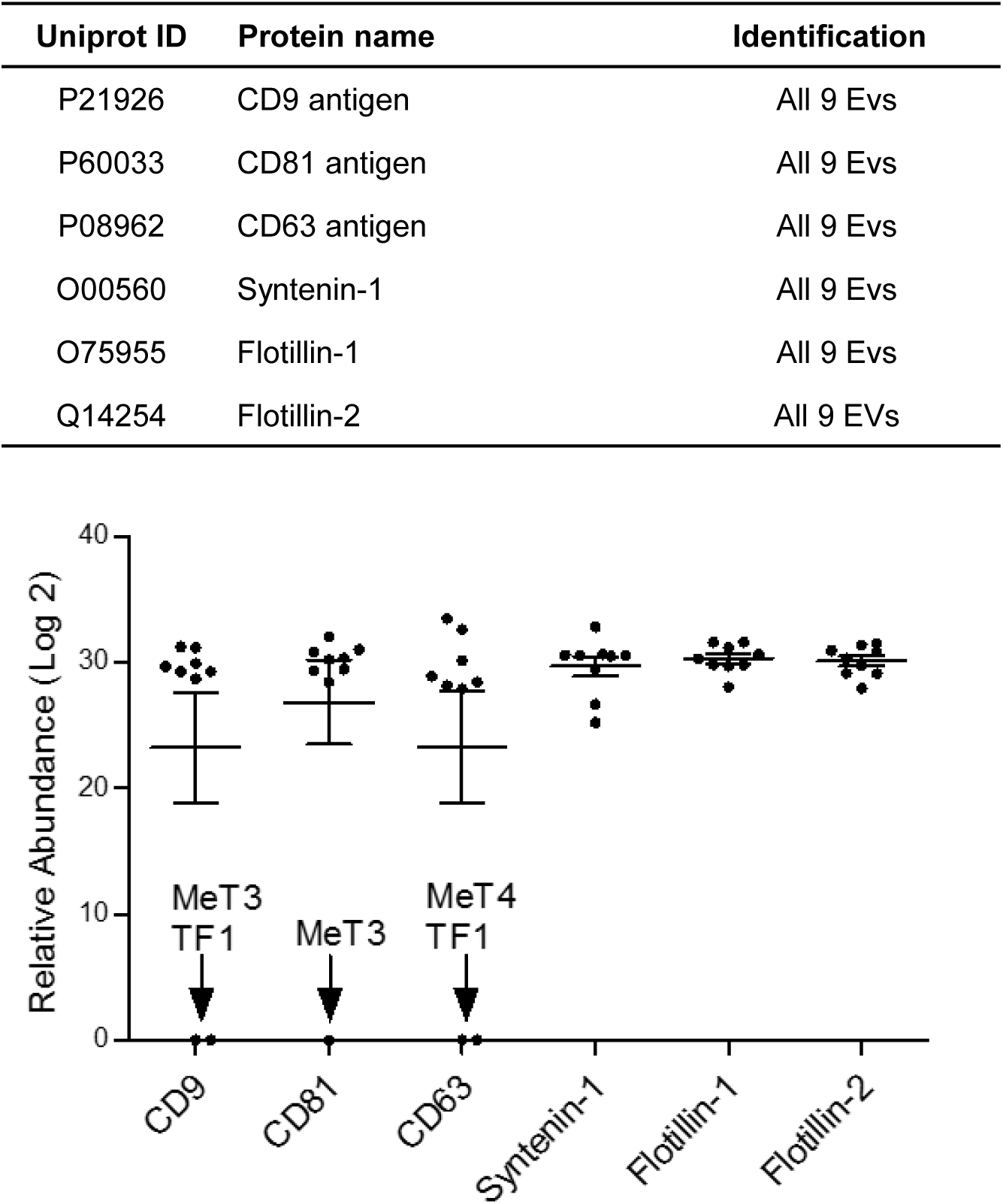
EV marker identification. Classical EV membrane marker proteins were identified in all five EV samples with similar abundance.

**Fig. S4.**
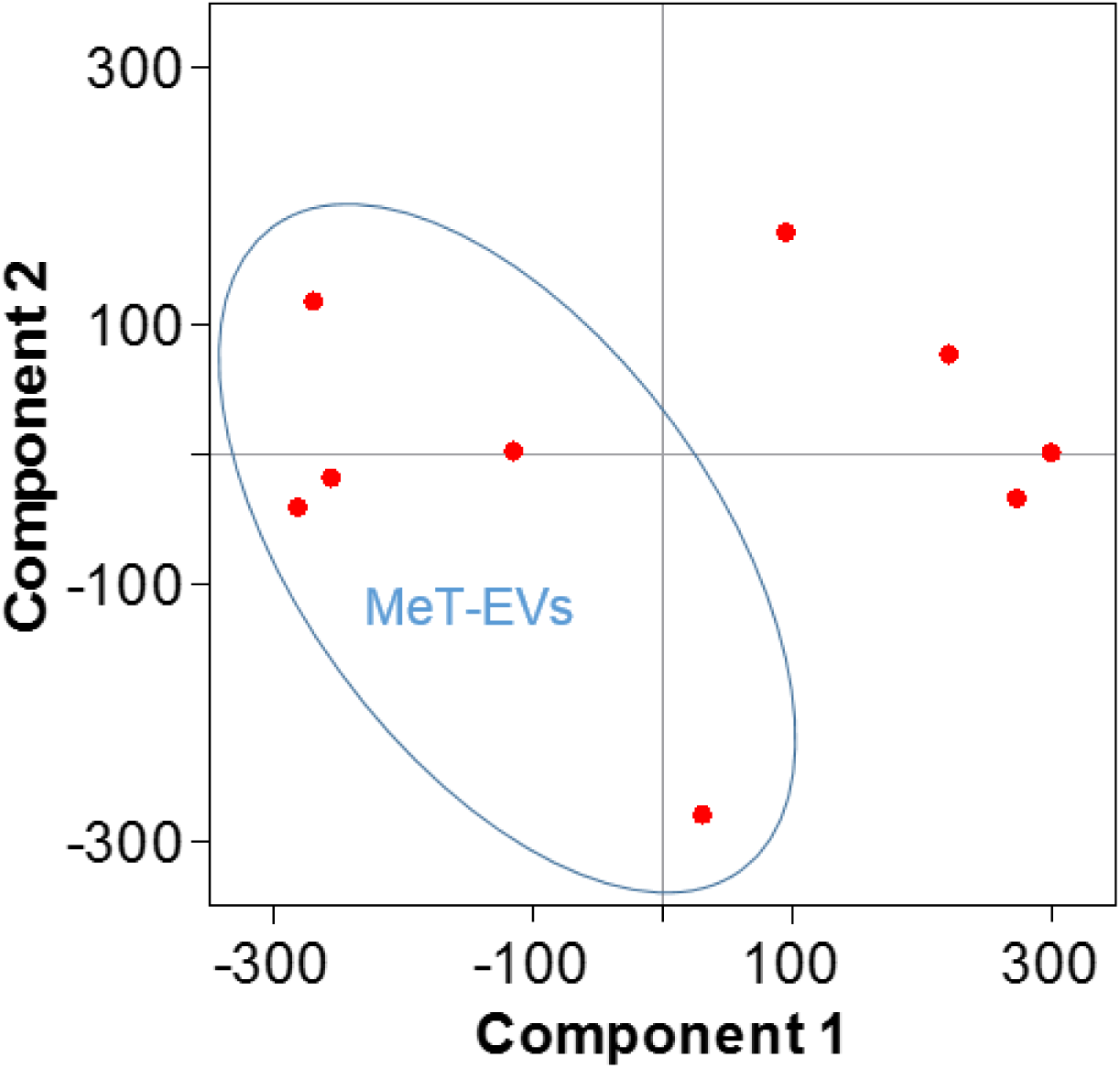
Principal component analysis of 9 EV proteome.

**Fig. S5.**
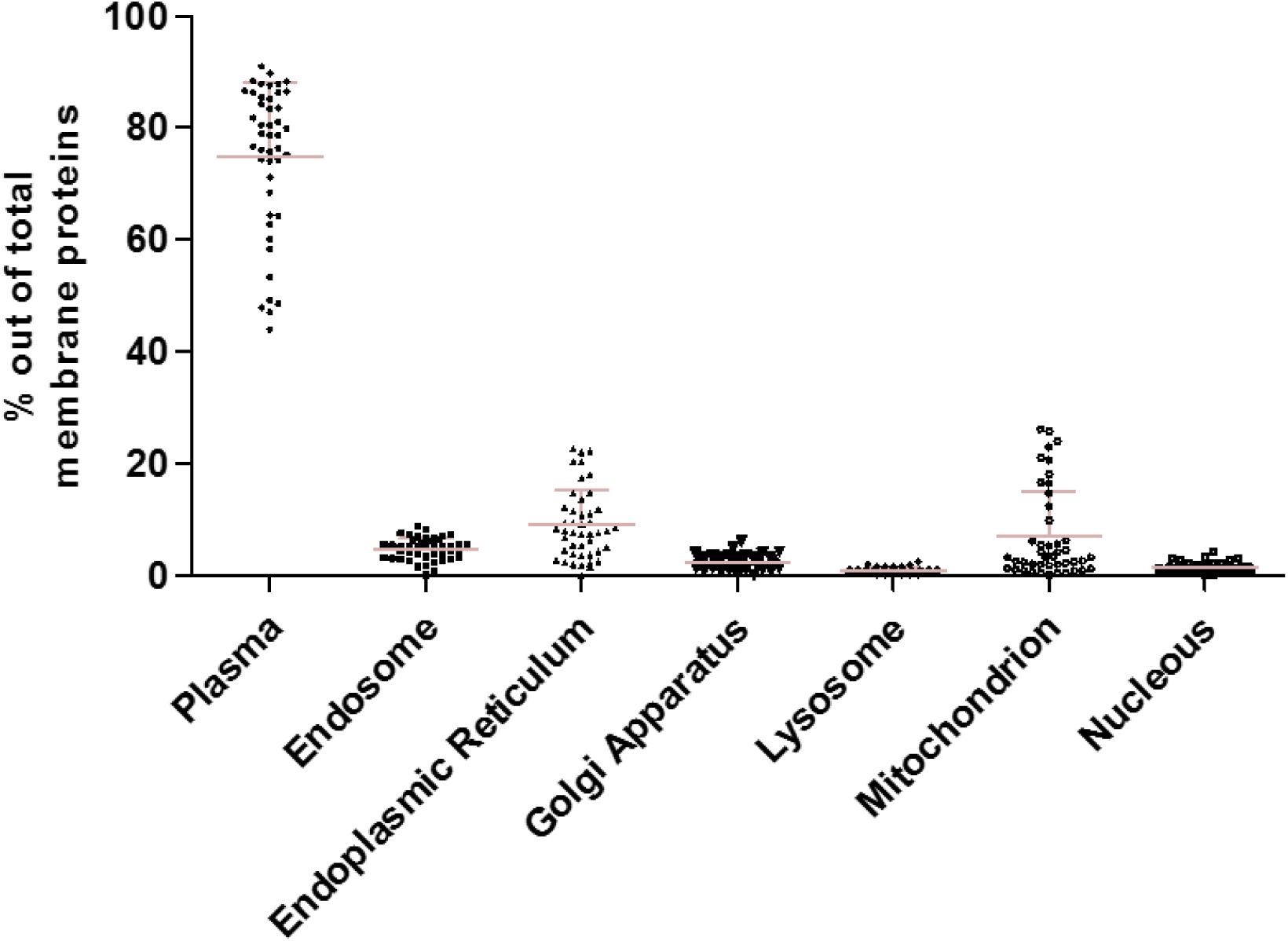
Proportion of organelle membrane proteins. The percentages of each origin of membrane proteins were calculated from 43 datasets in EVpedia. Localization was obtained from Uniprot and primary localization was used for analysis.

**Fig. S6.**
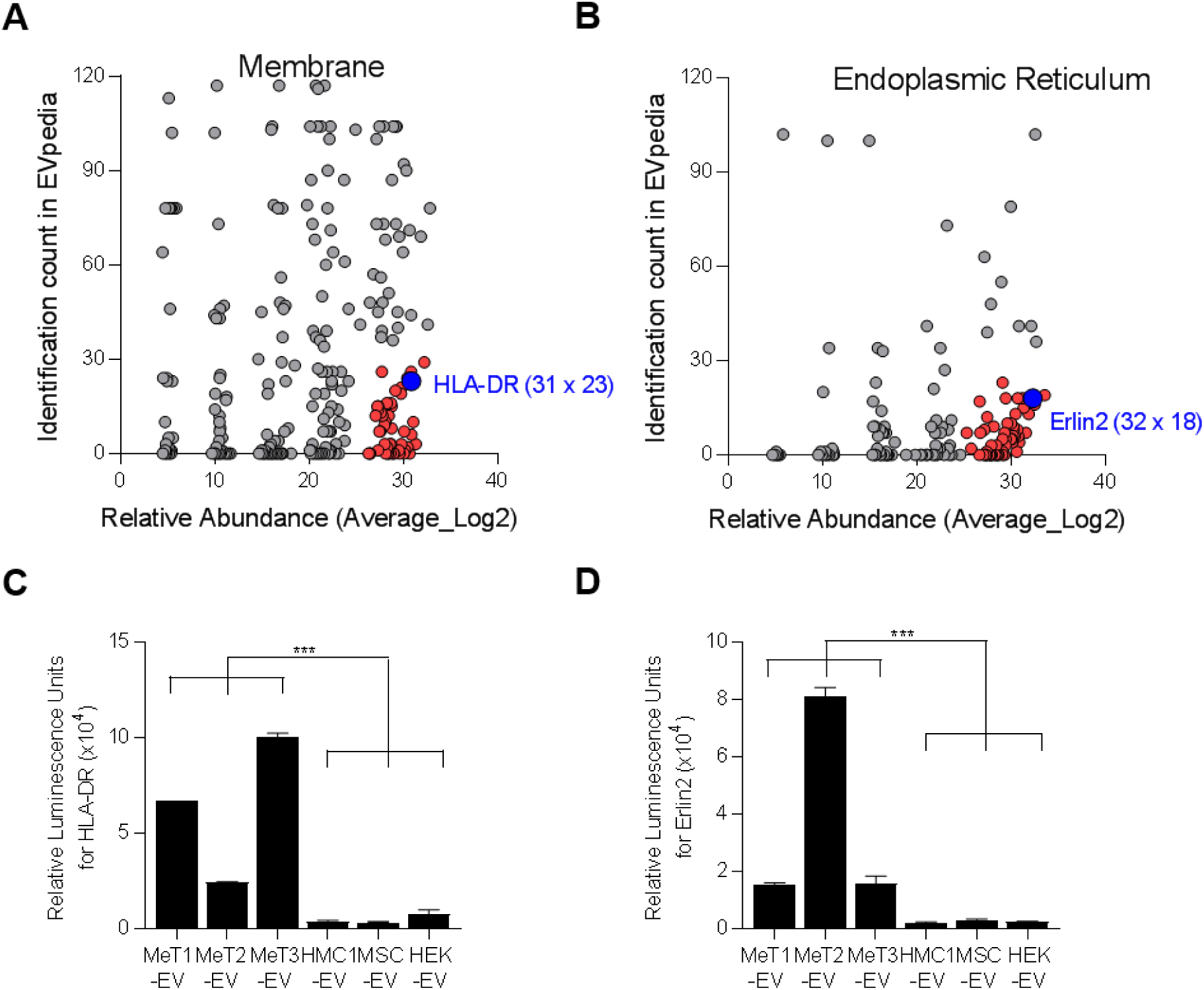
Validation of proteins by ELISA. Plasma membrane proteins (*A*) and endoplasmic reticulum membrane proteins (*B*) were plotted with their relative abundance from a mass spectrometry analysis and their identification count from the EVpedia database. Blue color are the final candidates for validation. IC; identification count. HLA-DR (*C*) and Erlin2 (*D*) were experimentally validated with direct ELISA. Data are presented as the mean ± SD. \*\*\**p* < 0.001.

**Fig. S7.**
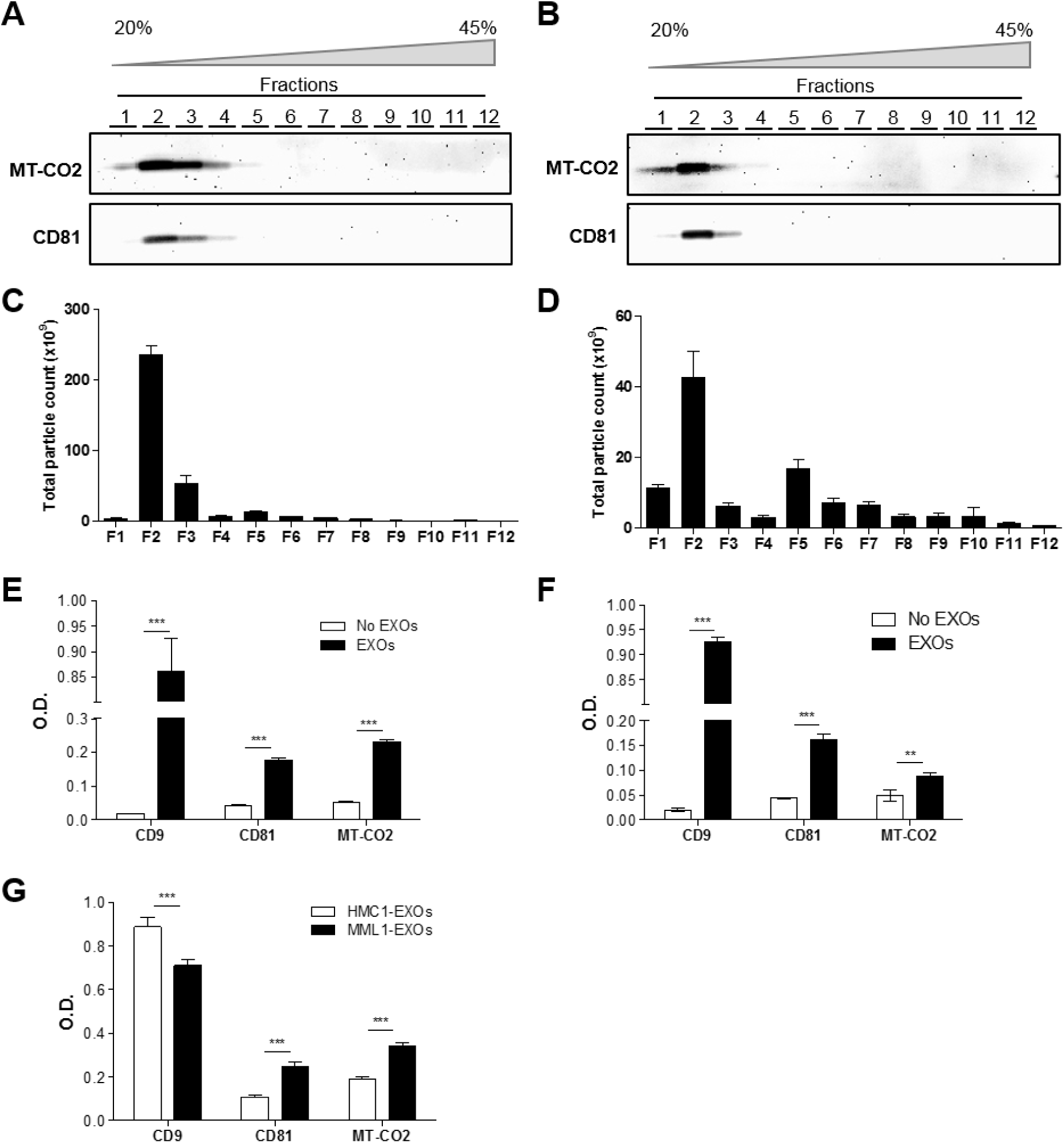
Mitochondrial MT-CO2 protein on the surface of EXOs. EXOs derived from MML1 (*A*) and HMC1 (*B*) were subjected to OptiPrep density gradient purification. MT-CO2 and CD81 expression in each fraction were visualized by Western blot. Particle numbers for each fraction from MML1-EXOs (*C*) and HMC1-EXOs (*D*) were determined by nanoparticle tracking analysis. Surface expression level of CD9, CD81, and MT-CO2 on MML1-EXOs (*E*) and HMC1-EXOs (*F*) was measured by direct ELISA. (*G*) The surface expression level was compared between MML1-EXOs and HMC1-EXOs. Data are presented as the mean ± SD. \*\**p* < 0.01, \*\*\**p* < 0.001.

**Fig. S8.**
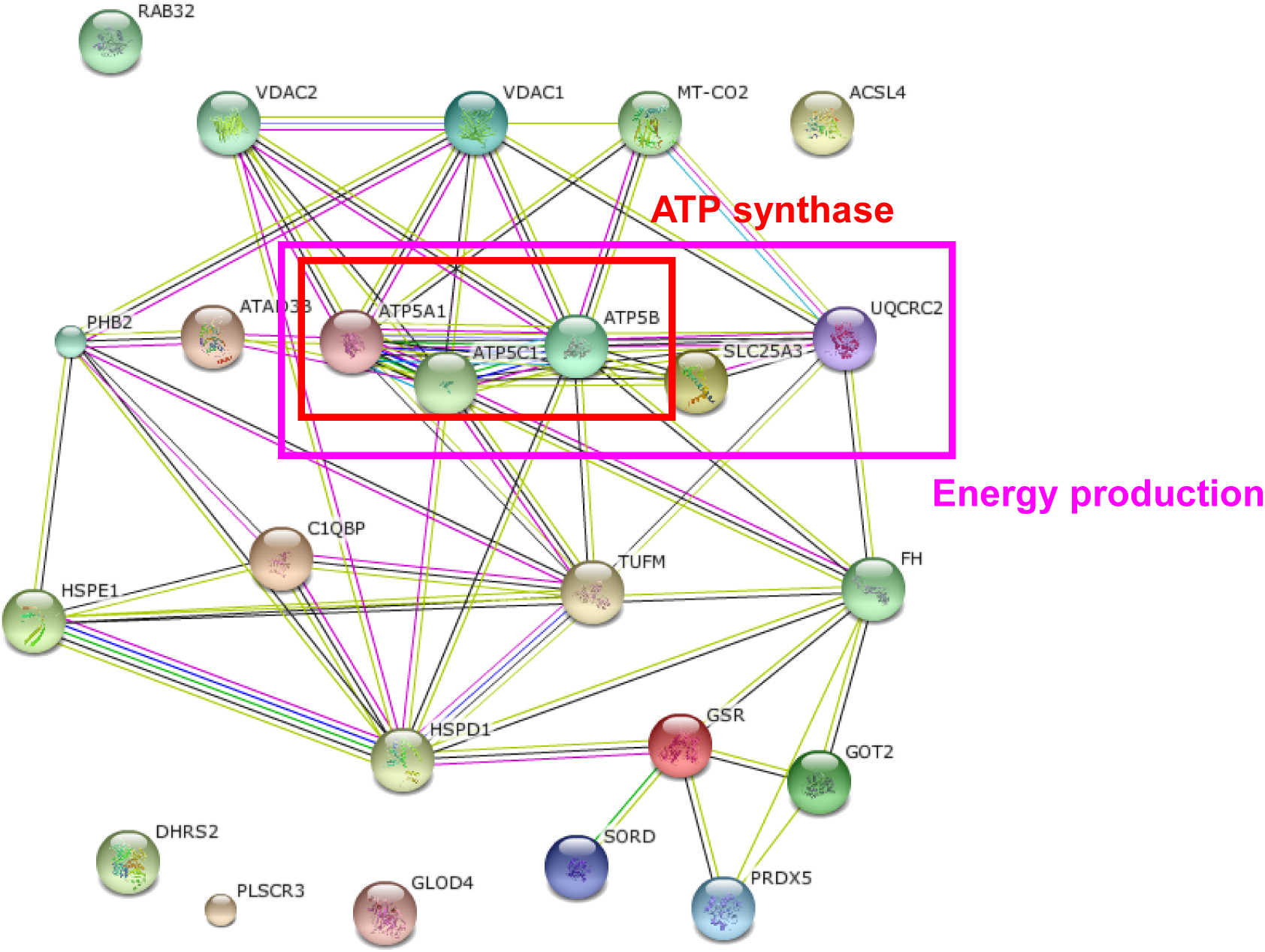
Protein-protein interaction network of mitochondrial proteins on MTCO2-EXOs. Identified mitochondrial proteins, including ATP synthase subunits and energy production machinery, interacted with each other.

**Fig. S9.**
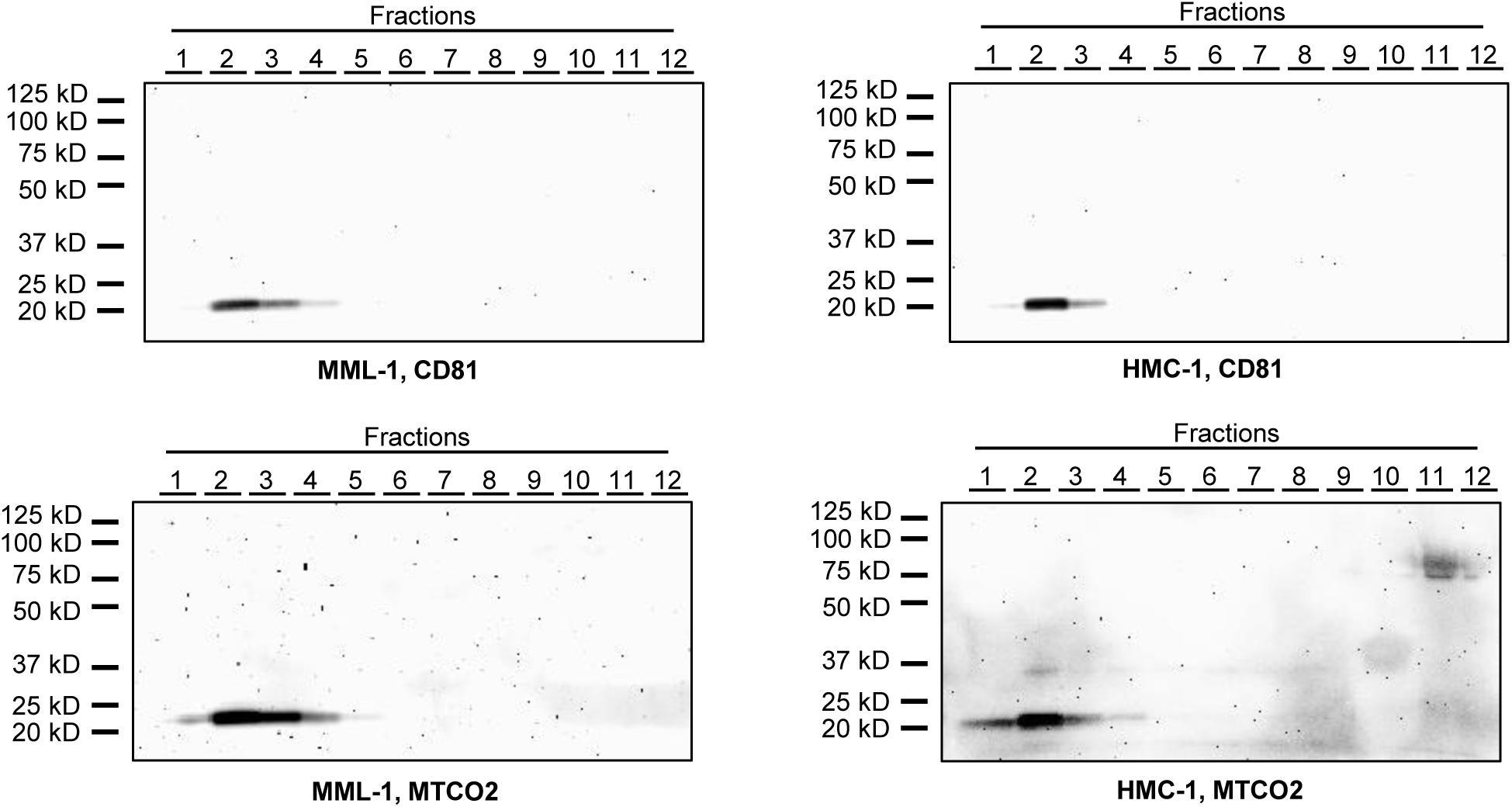
Full length images of Western Blot.

